# Meta-analysis reveals a distinct and uniform gut microbial signature associated with endocrine-disrupting chemicals-induced diabetes

**DOI:** 10.64898/2026.05.13.724769

**Authors:** Karthika Durairaj, Buvaneswari Gajendhran, Gowtham Manivel, Hariharan Gnanam, Krishnan Swaminathan, Mithieux Gilles, Ganesan Velmurugan

## Abstract

In recent years, the synergistic role of endocrine-disrupting chemicals (EDCs) and gut microbiota in the development of diabetes has been increasingly documented in rodent models. However, most studies have focused on one or two EDCs with varying doses and exposure durations, limiting the identification of a shared microbial signature associated with EDC-induced glucose dysregulation. This meta-analysis aimed to identify a common gut microbiome pattern across rodent studies involving diverse EDC exposures linked to glucose dyshomeostasis. A systematic search yielded 3,748 studies, of which ten met the inclusion criteria, comprising sequence data from 189 samples. These studies evaluated gut microbiota alterations in diabetes induced by various EDCs, including pesticides, food additives, and heavy metals, across different exposure conditions. Meta-analysis revealed a consistent reduction in microbial diversity and an increased Firmicutes/Bacteroidetes ratio following EDC exposure. At the phylum level, Firmicutes, Proteobacteria, Desulfobacterota, and Patescibacteria were significantly enriched. Although beneficial genera such as *Lactobacillus, Bifidobacterium*, and *Akkermansia* showed a decreasing trend, these changes were not statistically significant. In contrast, xenobiotic-associated genera including *Desulfovibrio, Pseudomonas, Parasutterella*, and *Candidatus Saccharimonas* were significantly increased. Notably, sulfate-reducing bacteria were the only inflammation-associated group consistently elevated. These microbial alterations were distinct from those observed in high-fat diet-induced diabetic models. This study identifies a distinct gut microbiome signature associated with EDC exposure in rodent models of glucose imbalance. These findings suggest unique microbiome-mediated pathways in EDC-induced diabetes and highlight potential microbial targets for early intervention in environmentally driven metabolic disorders.

## Introduction

The growing incidence of diabetes is attributed to both modifiable and non-modifiable risk factors. While sedentary lifestyles, unhealthy diets, obesity, and genetic predisposition are well-established contributors, recent attention has shifted toward the role of environmental pollutants and chemical exposures in the pathogenesis of diabetes (Neel & Sargis, 2011; Velmurugan *et al*. 2017a). Concurrently with the escalation of diabetes prevalence, the world has witnessed a massive production and release of toxic chemicals into the environment through industries, chemical-based agriculture and food production, and electronic wastes (Velmurugan *et al*. 2017a). A growing body of epidemiological and experimental research has identified an association between exposure to environmental chemicals and the incidence of metabolic diseases, particularly type 2 diabetes (Kuo *et al*. 2015; Velmurugan *et al*. 2020). These chemicals include a wide variety of compounds such as persistent organic pollutants pesticides, heavy metals, plasticizers, polycyclic aromatic hydrocarbons, dioxins and food additives that interfere with the endocrine system by altering the production, release, transport and action of hormones; these are termed as endocrine-disrupting chemicals (EDCs) (Diamanti-Kandarakis *et al*. 2009; La Merrill *et al*. 2020). Oral exposure to particularly *via* the food chain is the major pathway for the entry of EDCs into the human body. The gut microbiota comprises trillions of bacteria, fungi, and viruses and is recognized as a key player in the microbial metabolism of dietary compounds, drugs and EDCs (Carmondy & Turnbaugh, 2014; Spanogiannopoulos *et al*. 2016). Importantly, emerging research has demonstrated that EDCs can alter the gut microbial ecosystem, leading to dysbiosis that contributes to the pathogenesis of diabetes (Velmurugan *et al*. 2017a; Aguilera *et al*. 2020). The influence of gut microbiota on host glucose metabolism is mediated through various mechanisms, including modulation of bile acid profiles, production of microbial metabolites such as short-chain fatty acids (SCFAs) and branched-chain amino acids (BCAAs), and regulation of incretin hormones such as glucagon-like peptide-1 (GLP-1) (Li *et al*. 2023).

Faecal microbiota transplantation (FMT) studies in rodents have provided strong evidence supporting the causal role of gut microbiota in mediating EDCs-induced metabolic phenotypes (Velmurugan *et al*. 2017b; Hannsenn *et al*. 2021). The composition and metabolic activity of the gut microbiota may determine an individual’s susceptibility to the diabetogenic effects of environmental exposures, suggesting a personalized aspect of disease risk based on microbial profiles. Though there are several studies on rodent models reporting the changes in gut microbiota during EDCs-induced diabetes. But all these studies employed different animal models and the dose of EDCs used are largely not a realistic doses equivalent to human exposure. A common gut microbial signature associated with EDCs-induced diabetes and its variation with diabetes of other etiology especially high fat diet is not available.

## Methods and materials

### Study selection for meta-analysis

A comprehensive search term was entered into PubMed and the NCBI Sequence Read Archive (SRA) to retrieve an unbiased selection of studies investigating the effects of endocrine-disrupting chemicals on hyperglycemia and their impact on the murine gut microbiome. A total of 51 EDCs were employed as keywords alongside “gut microbiome” or “gut microbiota” (e.g: ‘‘organophosphates’’ [All Fields] AND ‘‘microbiome’’ [All Fields] OR ‘‘organophosphates’’ [All Fields] AND ‘‘microbiota” [All Fields]). The Systematic PubMed Search for Peer-Reviewed Articles were represented on Fig.S1. PubMed and the NCBI Sequence Read Archive (SRA) were used in a thorough search to find pertinent research investigating the effects of endocrine-disrupting chemicals (EDCs) on hyperglycemia and the mouse gut flora. Four independent researchers, blinded to each other’s decisions, independently reviewed the keywords and excluded irrelevant literature. Discrepancies were resolved through a consensus or the decision of a one author (Dr. G. Velmurugan). Our retrospective study adhered to the Preferred Reporting Items for Systematic Reviews and Meta-Analysis (PRISMA) guidelines (Fig.S2). All the search results were compiled and subjected to a structured screening process. Initially, the titles and abstracts of each article were reviewed, and their objectives were recorded in an Excel sheet to assist in identifying studies relevant to the research question. Duplicate records were removed, followed by the application of strict inclusion and exclusion criteria. The studies with reviews, meta-analysis or systematic reviews, clinical trials and the studies with human subjects, non-mammalian species, plants or in vitro models and studies on pregnant, neonatal, weaning and elderly murine models were excluded. Selected articles without publicly available metagenomics sequencing data were not included in the further analysis. Furthermore, studies that did not report gut microbial diversity using 16S rRNA gene or metagenomics sequencing, or those lacking publicly available raw sequencing data and metadata, were omitted. To be considered for inclusion, studies had to investigate the effects of endocrine-disrupting chemicals in murine models and provide evidence of EDC-induced hyperglycemia, indicated by elevated blood glucose levels. Preference was given to studies with accessible metagenomics or 16S rRNA sequencing data related to gut microbiota. Three independent reviewers conducted blinded evaluations of all articles based on these criteria, and the final list of eligible studies was curated through consensus. In cases of disagreement, discussions were held to reach a resolution. Reference lists of the included articles were also examined to ensure the comprehensiveness of the selection process. The overall article screening and selection workflow was illustrated in the PRISMA flow diagram, which details the number of studies identified, screened, excluded, and ultimately included in the final analysis. The data extraction was carried out using a pre-tested, standardized extraction form that captured several critical parameters relevant to the review. Key study characteristics, including authorship, year of publication, sample size, study design, duration, type of endocrine-disrupting chemical (EDC), and dosage, were systematically recorded. Detailed information on the animal model such as age and strain of the mice or rats - was also documented. Metabolic outcomes, particularly changes in fasting glucose and insulin levels, were extracted where available. In addition, comprehensive details related to the metagenomics data were collected, including sample type, DNA extraction kit used, and the sequencing platform employed for 16S rRNA gene analysis. Data were extracted from all available sources within each article, including the main text, tables, figures, supplementary materials, and appendices. Relevant data points were retained for potential inclusion in the meta-analysis, depending on their completeness and consistency with the review criteria.

### Data collection and sequence retrieval

A total of 10 published studies were selected based on the availability of raw sequencing data related to gut microbiota profiling in relation to metabolic or environmental exposures. The datasets included both 16S rRNA gene amplicon sequencing and whole metagenome shotgun sequencing data. Raw sequencing reads were downloaded from publicly available repositories such as the NCBI Sequence Read Archive (SRA) using accession numbers provided in the respective publications. The datasets encompassed a mixture of single-end and paired-end reads, generated using various sequencing platforms (e.g., Illumina MiSeq, HiSeq).

### Sequence Processing and QIIME 2 Analysis

All downloaded raw reads were processed using **QIIME 2** (version 2024.10.1) for standardized analysis (Bolyen *et al*., 2019). For 16S rRNA data, initial demultiplexing and quality assessment were performed. Single-end and paired-end reads were separately processed using the **DADA2 plugin** for quality filtering, denoising, chimera removal, and feature table construction. Taxonomic classification was performed using a pretrained Greengenes classifier based on the 99% identity threshold, targeting the V3–V4 region of the 16S rRNA gene. For shotgun sequence data, appropriate preprocessing (host removal, quality filtering) was performed prior to downstream taxonomic profiling. All datasets were normalized and aggregated at various taxonomic levels (phylum to genus) for comparative analysis. Metadata was curated uniformly across all studies to enable group-wise comparisons and diversity analyses.

### Statistical analysis

Meta-analyses for four alpha diversity measures were conducted using the R packages meta and metafor (R version 4.1.0) (Balduzzi *et al*., 2019). Random-effects models were applied, with effect sizes calculated as Hedges’ g standardized mean differences (SMDs). Results were considered statistically significant when the 95% confidence interval (CI) did not include zero. Heterogeneity across studies was assessed using the I^2^ statistic (percentage of total variation due to heterogeneity), the between-study variance (τ^2^), and Cochran’s Q test p-value. To standardize alpha diversity indices across studies, values were mean-centered to zero and scaled to unit variance. Boxplots were generated in GraphPad Prism (version 8.3.0). For taxonomic analyses, bacterial counts were converted to relative abundances. Taxa were retained only if they had an average relative abundance ≥0.005% and were present in ≥5% of samples within a study. Random-effects meta-analysis models with inverse variance weighting and the Der Simonian– Laird estimator were used to pool adjusted estimates and their standard errors. Analyses were restricted to taxa for which effect estimates and standard errors were available in at least 50% of included studies. Sensitivity analyses were performed by sequentially omitting one study at a time to evaluate the robustness of the findings. All statistical tests were two-sided, and p-values <0.05 were considered statistically significant after adjustment for multiple testing.

## Results

### Study selection and characteristics

A total of 51 EDCs were employed as keywords alongside “gut microbiome” or “gut microbiota” (e.g: ‘‘organophosphates’’ [All Fields] AND ‘‘microbiome’’ [All Fields] OR ‘‘organophosphates’’ [All Fields] AND ‘‘microbiota’’ [All Fields]). Our retrospective study adhered to the Preferred Reporting Items for Systematic Reviews and Meta-Analysis (PRISMA) guidelines. A total of 1330 unique studies were identified through our search methodology. To ensure meticulousness in study selection, four members of our laboratory, specializing in the microbiome field and well-versed in relevant terminology and methodologies, redundantly reviewed these studies to assess their study design and methodological relevance for potential inclusion. The detailed flowchart depicted in conforms to PRISMA guidelines, providing an overview of the search results. Our literature search encompassed 102 diverse combinations of keywords, resulting in the retrieval of 1330 articles. However, after applying the exclusion criteria, 1323 papers were deemed incongruent with the objectives of our study. Subsequently, a comprehensive review was conducted on high-throughput 16S rRNA sequence data sourced from 10 distinct studies, scrutinizing their full-text content for potential inclusion in further meta-analysis (Fig. 1).

**Fig. 1.**
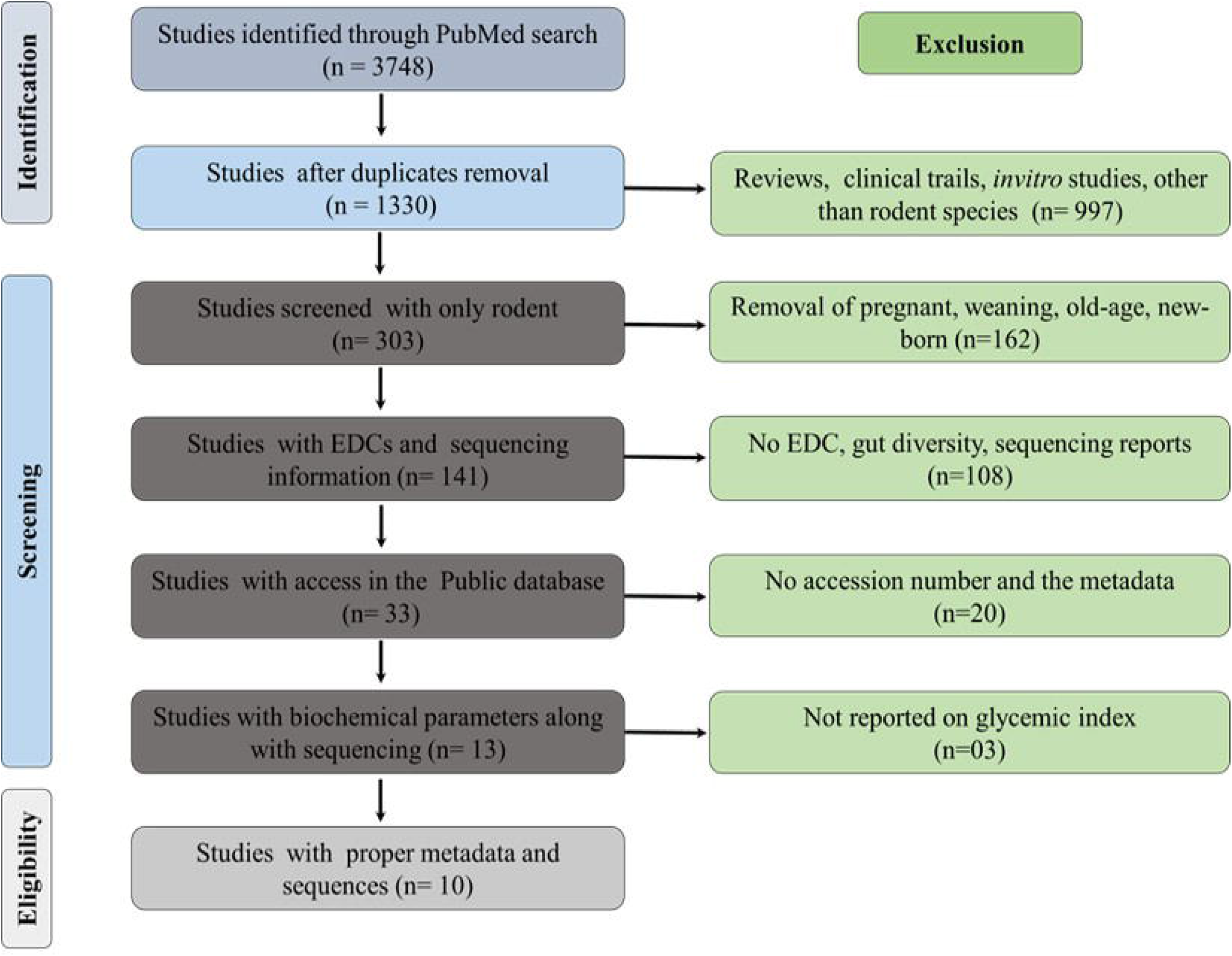
PRISMA flow diagram for the meta-analysis. A schematic representation of the meta-analysis workflow for microbiome sequencing datasets associated with endocrine-disrupting chemicals (EDCs) and hyperglycaemia

Finally, according to the inclusion criteria, 10 articles were reviewed for their full-text content and included for further meta-analysis (Table 1) (Suez *et al*., 2014; Chassiang *et al*., 2015; Wang QP *et al*., 2018; Ruan Y *et al*., 2019; Liang Y *et al*., 2019; Zhan J *et al*., 2019; Min *et al*., 2020; Wang J *et al*., 2021; Liu T *et al*., 2020; Mesnage *et al*.,2021; Shi *et al*.,2021). There is no sex bias in choosing the animal model across all studies. Out of the 10 included studies, a total of 14 different EDC treatments were investigated. These comprised food additives (n = 6 studies), heavy metals (n = 5 studies), and pesticides (n = 3 studies), in relation to the progression of type 2 diabetes mellitus and gut microbiota (Fig. S3A). The majority of studies used C57BL/6 mice as the rodent model, while a few employed Sprague–Dawley rats and Kunming mice (Fig. S1B). For microbiome sequencing, most studies targeted the V4 region of the 16S rRNA gene and used the Illumina MiSeq platform (Fig. S3C–D).

Consistent with the inclusion criteria, all selected studies reported an increase in blood glucose levels along with other biochemical alterations, including elevated total cholesterol and lipid profile across most studies. Upon EDC administration, increased activities of AST, ALT, and ALP enzymes were observed in five studies. Notably, six studies assessed faecal SCFAs, with two of these also measuring serum SCFAs (Table S1). Information on microbiome dataset availability, including accession numbers and the list of samples from each study used for further analysis, is provided in Table S2.

### Effect of EDC exposure on microbial diversity

We assessed alpha diversity using three common metrics-Chao1, Shannon index, and Simpson index across the selected studies. To account for varied baseline states, values were scaled to the geometric mean of EDC-exposed samples within each study. Visual inspection indicated no consistent alteration in alpha diversity, with considerable heterogeneity in the direction of effects. For the Chao1 index, the pooled random-effects estimate indicated no significant difference in richness following EDC exposure (log2 fold change = 0.09, 95% CI: –0.86 to 1.04, p = 0.85). Substantial heterogeneity was observed (I^2^ = 93.95%, τ^2^ = 0.4074, H^2^ = 14.39). For the Shannon index, which reflects microbial diversity, no consistent effect of EDC exposure was detected (log2 fold change = –0.12, 95% CI: –0.96 to 0.72, p = 0.78), with significant heterogeneity (I^2^ = 74.63%, τ^2^ = 0.0413, H^2^ = 3.94). Similarly, the Simpson index, reflecting evenness, did not show a significant pooled effect (log2 fold change = –0.25, 95% CI: –1.11 to 0.61, p = 0.57), though heterogeneity was again high (I^2^ = 97.46%, τ^2^ = 0.4273, H^2^ = 39.47). EDC exposure does not consistently alter alpha diversity, although substantial heterogeneity exists across studies (Fig. 2A-C). The principal coordinates analysis (PCoA) based on Bray-Curti’s dissimilarity and Jaccard distance at the genus level revealed partial separation between EDC-exposed and control groups, indicating shifts in community composition. However, clustering patterns were not consistent across all studies, suggesting that the extent of EDC-induced microbial perturbations may depend on factors such as exposure type, dose, and host background. Both Bray–Curtis and Jaccard metrics demonstrated substantial inter-study variation but still supported the presence of distinct compositional differences attributable to EDC exposure. (Fig. 3A-B).

**Fig. 2.**
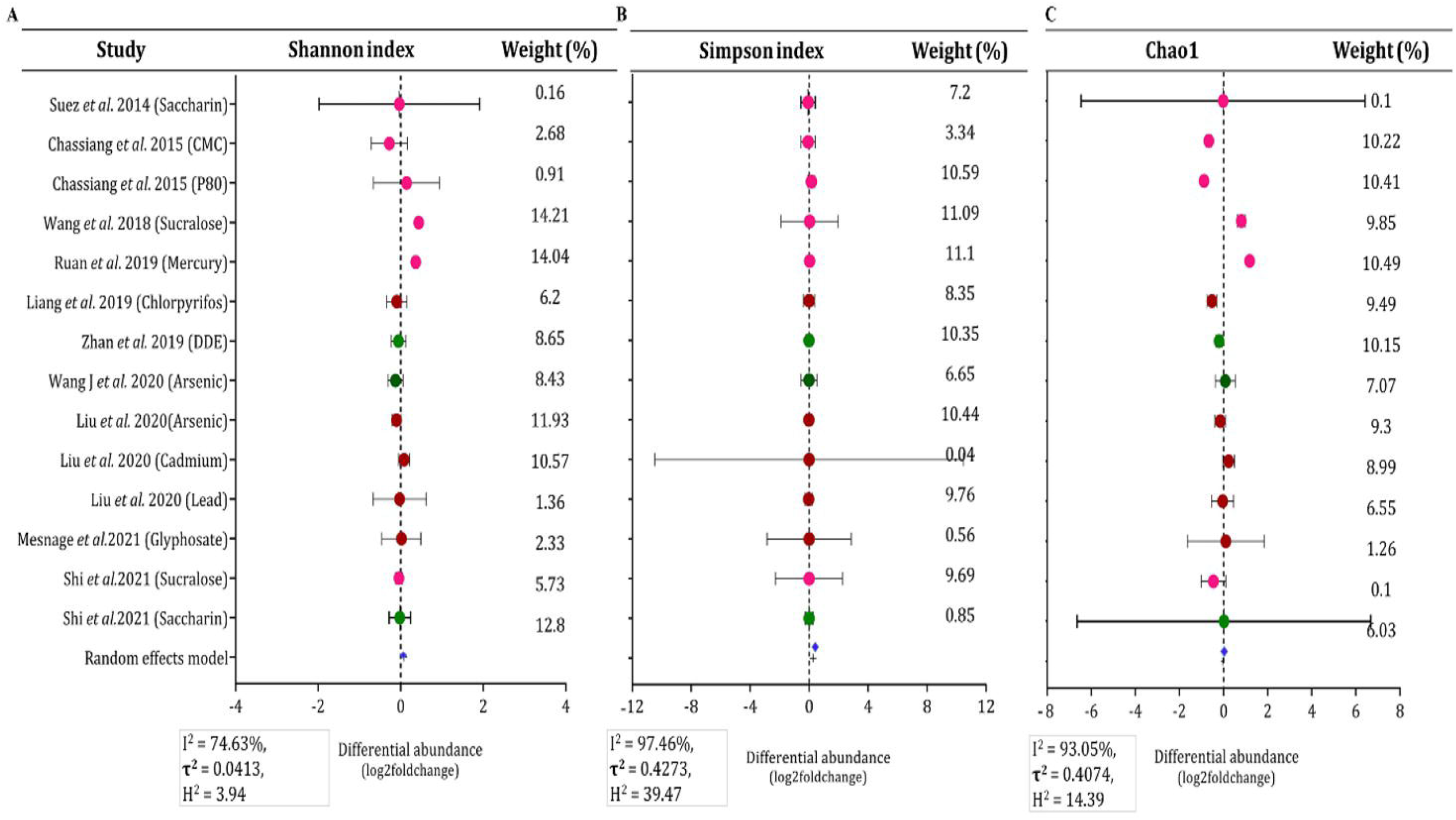
Forest plots showing the effects of various chemical exposures on microbial diversity indices. A - Meta-analysis of studies investigating the impact of different chemicals on gut microbiota diversity using the Chao1 index. B - Shannon index. C - Simpson index.

**Fig. 3.**
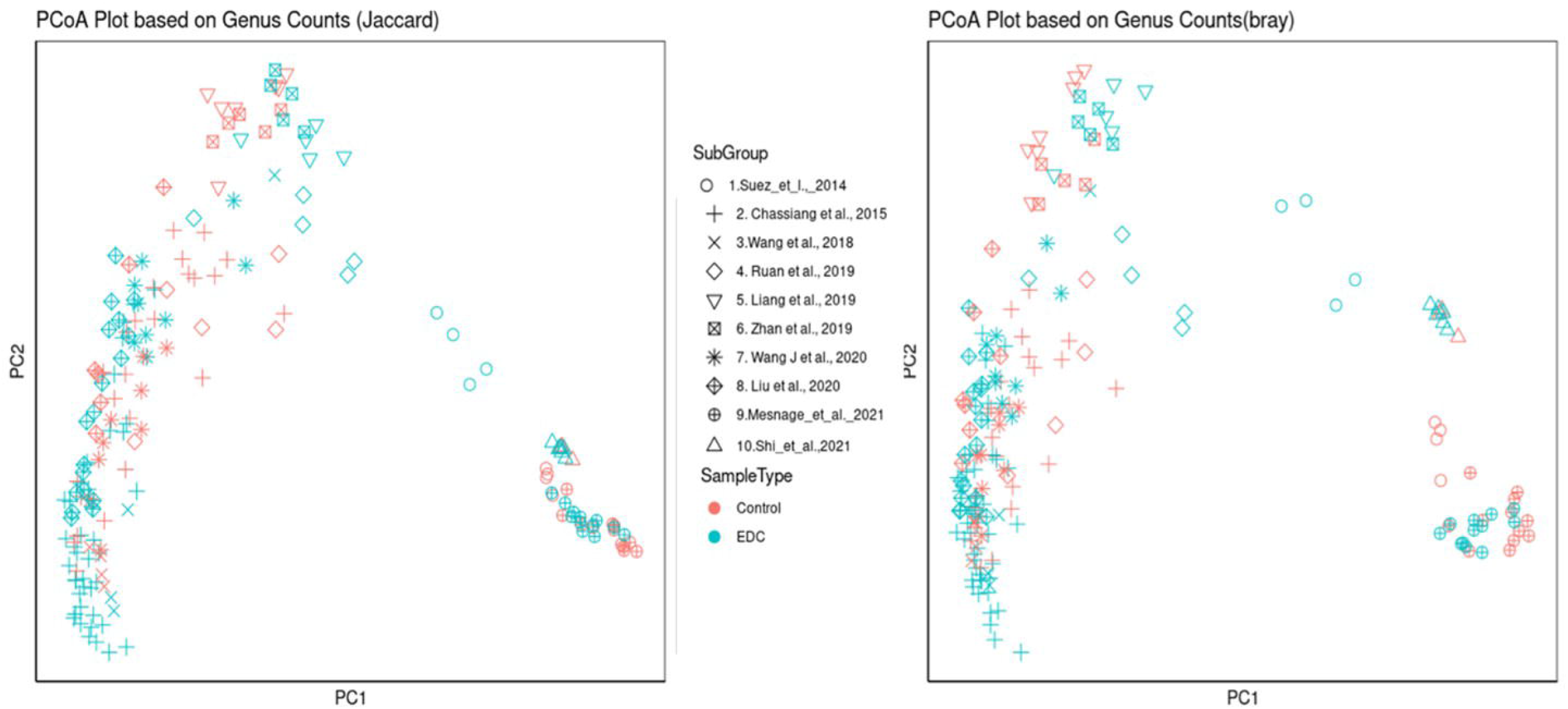
Principal Coordinates Analysis (PCoA) of Genus-Level Microbial Profiles. A - PCoA plots were generated based on genus-level counts using Jaccard. B - Plots using Bray–Curti’s distance metrics

### EDCs alter the microbial composition on phyla level

The relative abundance at the phylum level showed a major composition of *Bacteroidetes* and *Firmicutes*, with minor contributions *from Actinobacteria, Proteobacteria, and Desulfobacterota*, and trace amounts of *Verrucomicrobiota and Patescibacteria* on EDCs (Fig. 4A) and the individual EDCs categories (Fig. S4). The weight estimation of individual phyla level was calculated with Log2 foldchange values. An alteration in the *Firmicutes/Bacteroidetes* (F/B) ratio was observed in the EDC treatment group compared to controls (Fig. 4B). Heatmaps revealed *that Patescibacteria and Desulfobacterota* were significantly increased across all studies (Fig. 4C), while *Bacteroidot*a consistently decreased among the selected studies. The pooled analysis at the phylum level across all studies demonstrated a significant pattern associated with EDC treatment (Fig.4D). The individual phyla levels among the studies were altered among based on the EDCs (Fig. 5A-H).

**Fig. 4.**
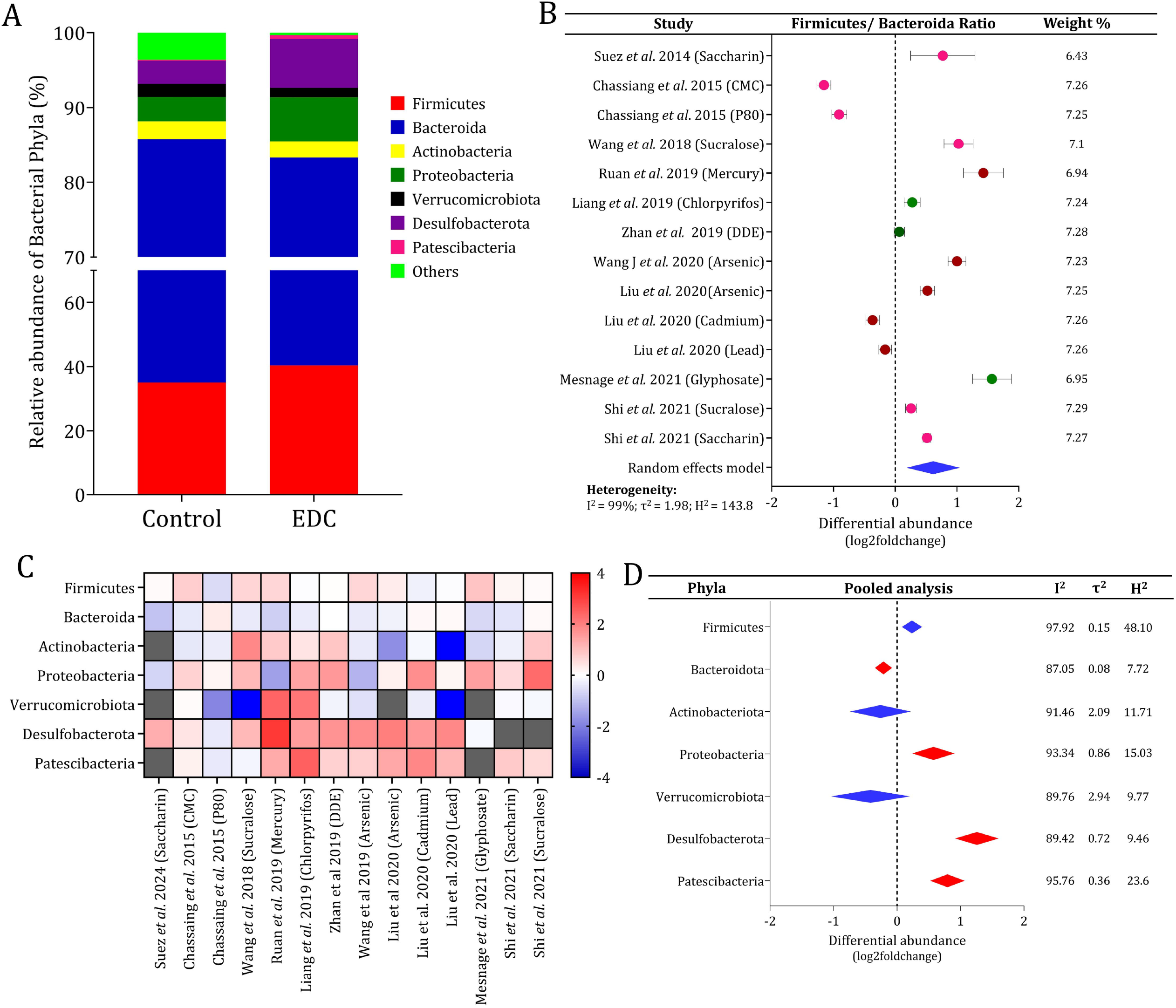
Endocrine-disrupting chemicals (EDCs) alter the gut microbiota composition at the phylum level. **(A)** Relative abundance of bacterial phyla in control and EDC-exposed groups.**(B)** Forest plot showing study-specific and pooled estimates of changes in the Firmicutes/Bacteroidetes ratio. **(C)** Heatmap depicting differential abundance (log2 fold change) of major bacterial phyla. **(D)** Pooled meta-analysis of phylum-level changes.

**Fig. 5.**
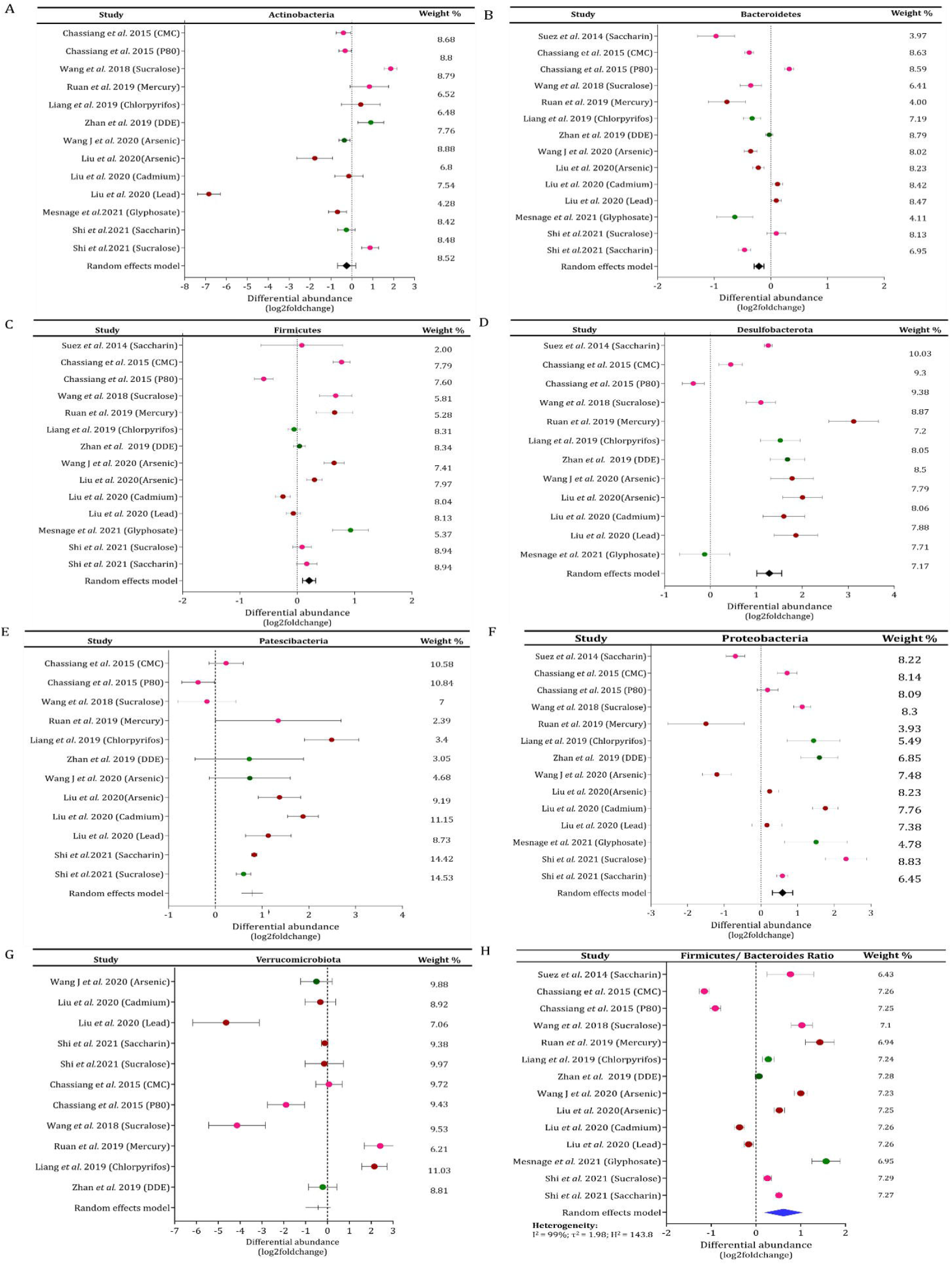
Individual phyla level observation for meta-analysis. *A) Actinobacteria, B) Bacteroidetes, C) Desulfobacterota, D) Firmicutes, E) Patescibacteria, F) Proteobacteria, G) Verrucomicrobiota, H) Firmicutes/Bacteroidetes ratio*

### Genera level alteration occurred in response to EDCs

For the genus-level analysis, only those genera that were reported in more than six of the ten selected studies were included to ensure consistency and reliability of the comparison. The shortlisted genera were subsequently categorized based on their predicted functional profiles generated through PICRUSt analysis. Accordingly, the genera were grouped into three major categories: beneficial taxa, xenobiotic-associated taxa, and inflammation-associated taxa. The EDC exposure resulted in widespread decreases in several beneficial genera. The heatmap indicated consistent negative fold-change patterns for *Akkermansia, Bifidobacterium, Lactobacillus*, and *Faecalibacterium* across multiple studies (Fig. 6A). The pooled analysis further supported this trend, showing an overall negative effect size for most beneficial taxa (Fig. 6B), suggesting that EDC exposure suppresses genera involved in maintaining gut barrier integrity, energy metabolism, and short-chain fatty acid production.

**Fig. 6.**
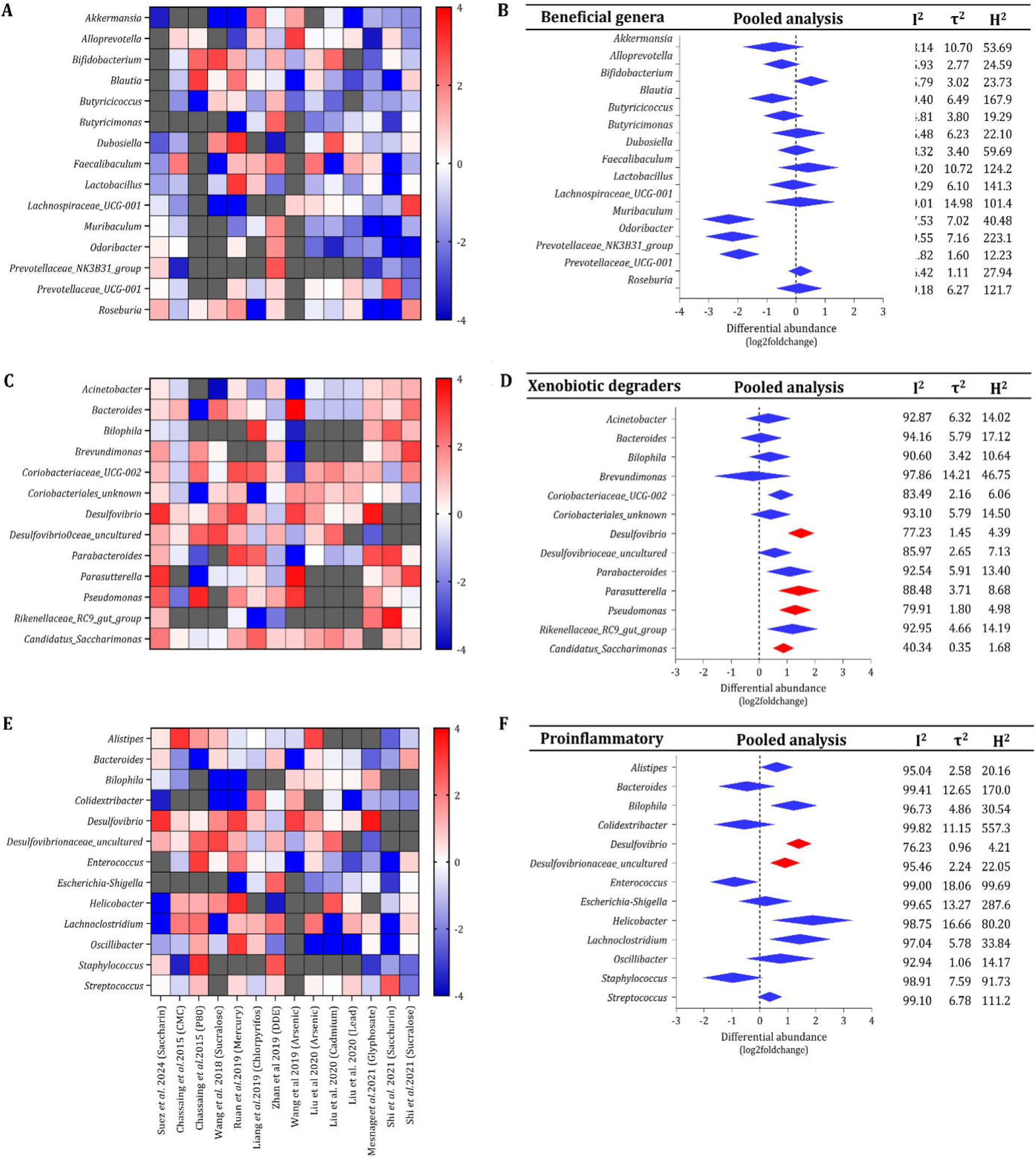
EDC exposure leads to alterations in the functional profiles predicted from genus-level microbiota data. **(A-B)** Heatmap and pooled analysis for the beneficial associated genera. **(C-D)** Heatmap and pooled analysis for the xenobiotic degraders associated genera. **(E-F)** Heatmap and pooled analysis for the proinflammation associated genera

Genera associated with xenobiotic degradation exhibited a predominantly positive response to EDC exposure. The heatmap demonstrated increased abundance patterns for *Bacteroides, Bilophila, Desulfovibrio, Parabacteroides*, and other xenobiotic-degrading genera (Fig. 6C). The pooled forest plots showed positive effect sizes for several of these taxa (Fig. 6D), indicating that EDCs may selectively enrich microbial groups capable of metabolizing environmental toxicants or participating in sulfur and bile acid metabolism. Genera categorized as proinflammatory displayed a mixed but generally elevated abundance following EDC exposure. The heatmap revealed increased representation of *Alistipes, Collinsella, Escherichia/Shigella, Staphylococcus*, and *Streptococcus* in multiple studies (Fig. 6E). The pooled effect-size estimates indicated a positive shift for many of these genera (Fig. 6F), suggesting that EDC exposure promotes the expansion of microbial communities linked to inflammation, gut permeability, and metabolic dysregulation.

### Differentially abundance of bacterial genera due to EDCs

The LEfSe analysis revealed distinct taxonomic signatures across the experimental groups, as illustrated in the cladogram. Several bacterial genera showed significant differential abundance, indicating group-specific microbial alterations in response to the treatments. Genera enriched in the treatment group were represented by colored branches (e.g., red, yellow, green, blue), whereas black nodes correspond to taxa that did not differ significantly between groups. The cladogram highlights clear clustering patterns, demonstrating that specific microbial lineages contributed strongly to the observed differences. Notably, treatment-enriched genera included *Streptococcus, Lactobacillus, Enterococcus, Dubosiella, Oscillibacter*, and *Butyricicoccus*, which were predominantly represented in the red-colored clades. In contrast, genera such as *Prevotellaceae UCG-001, Alloprevotella, Muribaculaceae*, and *Colidextribacter* were enriched in the control group, represented by blue branches. The hierarchical structure of the cladogram indicates that these shifts occurred across multiple taxonomic levels, emphasizing that EDC exposure induced distinct and consistent microbial alterations. These differentially abundant taxa collectively highlight the microbial signatures associated with each group and support the evidence of gut microbiota dysbiosis driven by the treatment (Fig. 7).

**Fig. 7.**
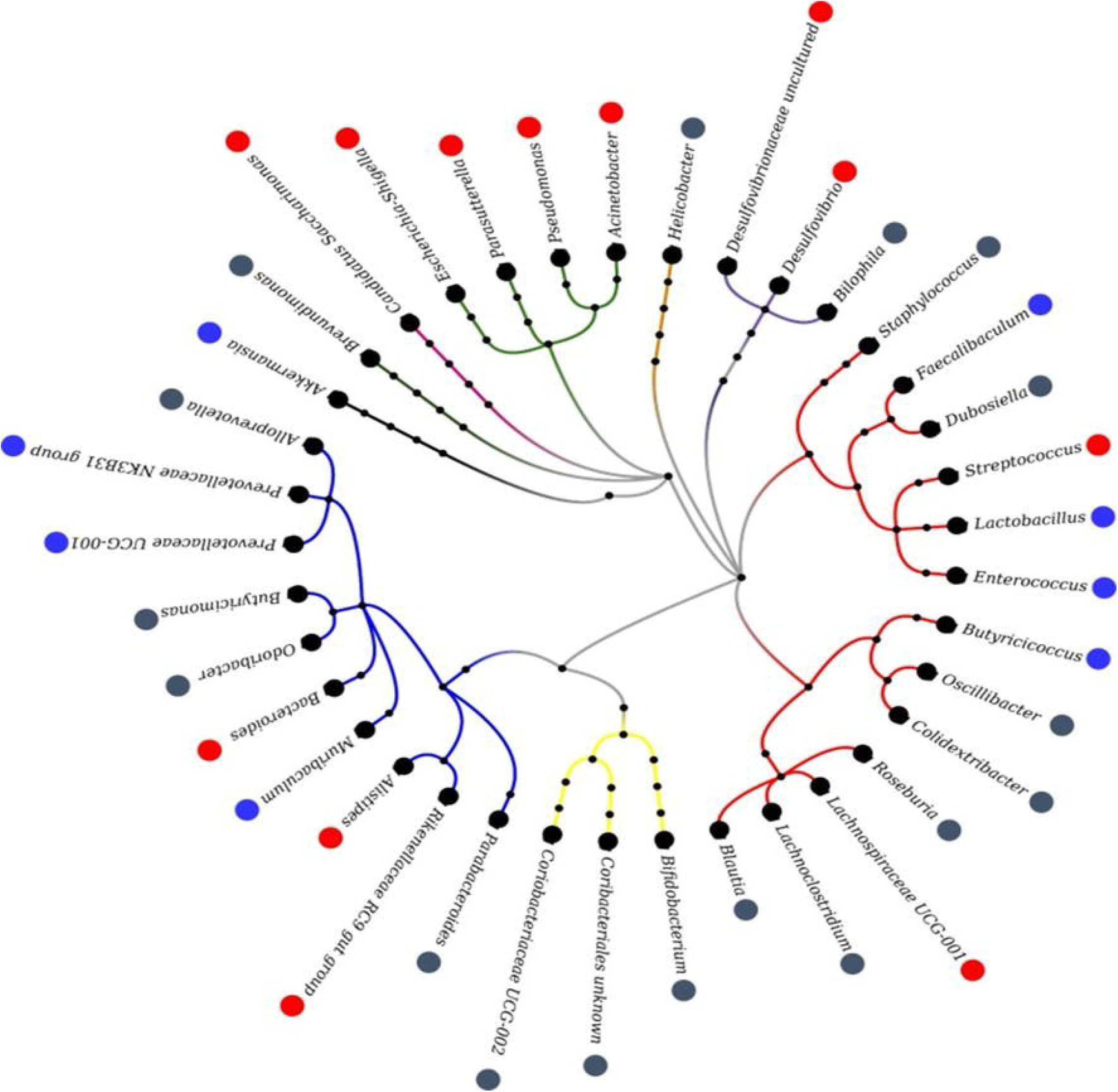
A Differentially abundant gut bacterial taxa identified by LEfSe analysis. Cladogram showing the taxonomic hierarchy of gut bacterial genera that differ significantly between study groups, as identified by LEfSe analysis (LDA score above the significance threshold). Colored branches and nodes represent taxa enriched in each group, while black nodes indicate taxa not significantly differentially abundant. B. Venn diagram representing the variation in gut microbiota among the EDC-induced and high fat diet-induced diabetes models.

## Discussion

The present meta-analysis synthesizes 16S rRNA sequencing data from 10 studies encompassing 14 distinct EDC exposures (food additives n=6, heavy metals n=5, pesticides n=3), revealing consistent gut microbiota dysbiosis in hyperglycemia models: phylum-level increases in *Firmicutes/Bacteroidetes (F/B) ratio, Desulfobacterota, and Patescibacteria* alongside *Bacteroidota* depletion; genus-level depletions of beneficial taxa (*Akkermansia, Bifidobacterium, Lactobacillus, Faecalibacterium*); and enrichments of xenobiotic-degraders (*Bacteroides, Bilophila, Desulfovibrio, Parabacteroides*) and pro-inflammatory genera (*Alistipes, Collinsella, Escherichia-Shigella, Streptococcus*). These shifts align with biochemical changes reported across studies, including hyperglycemia, dyslipidemia, and elevated ALT/AST/ALP, supporting EDC-induced T2DM progression via microbial mediation. The observed F/B ratio elevation echoes EDCs-diabetes meta-analyses, where first demonstrated higher *Firmicutes*-driven energy harvest in ob/ob mice (Turnbaugh *et al*., 2006; Fan & Pedersen, 2021) and confirmed this in human T2DM cohorts with reduced butyrate-producers. However, EDC-specific enrichments in *Desulfobacterota* (e.g., *Bilophila*) and Patescibacteria distinguish these profiles. Depleted *Akkermansia muciniphila*, a mucin-degrader producing propionate and strengthening barrier integrity, parallels findings in metformin-treated T2DM (Yan *et al*., 2021) and obesogenic diets (Everard *et al*., 2013), suggesting shared protective roles against metabolic endotoxemia. Alpha-diversity metrics showed no pooled changes (Chao1 log2FC 0.09, 95% CI −0.86–1.04, p=0.85; Shannon −0.12, p=0.78; Simpson −0.25, p=0.57), with high heterogeneity (I^2^ 74–97%), despite beta-diversity separation (PCoA Bray-Curtis/Jaccard). This compositional restructuring without richness loss contrasts uniform diversity declines in obesity meta-analyses (e.g., I^2^<50% for Shannon reductions), likely due to EDC class variability favoring resilient xenobiotics-degraders over broad suppression (Li *et al*., 2021). The gut microbiome serves as a key mediator in environment-health interactions, aiding in the detoxification of harmful compounds, and thus merits inclusion in risk assessment frameworks (De Filippis *et al*., 2024). PICRUSt-predicted functions reinforce this: suppressed SCFA pathways correlate with 6 studies reporting fecal/serum SCFAs. Mechanistically, these signatures promote T2DM via interconnected pathways. Xenobiotic-enrichments enhance EDC deconjugation (e.g., Bacteroides esterases), increasing circulating agonists for PPARγ/estrogen receptors in adipose/liver (Martyniu *et al*. 2022; Aguilera *et al*., 2020) Pro-inflammatory blooms (*Escherichia-Shigella, Streptococcus*) elevate LPS/TMAO, driving TLR4/NF-κB activation and metaflammation, akin to high-fat diet models (Cani *et al*., 2011). Beneficial taxa losses impair GLP-1 secretion and tight junctions, exacerbating glucotoxicity; LEfSe cladograms pinpointing *Oscillibacter/Butyricicoccus* gains versus *Prevotellaceae* losses further delineate EDC-specific clades. Compared to obesity, where SCFA deficits dominate, EDC dysbiosis uniquely amplifies chemical toxicodynamic, as human cohorts link urinary phthalates/Bisphenol-A to similar microbial shifts and HbA1c rises (Hinault *et al*., 2023). Study heterogeneity arises from models (C57BL/6 dominant, some Sprague-Dawley/Kunming), V4-region sequencing uniformity notwithstanding, exposure durations, and absent sex stratification despite no reported bias. Retrospective design limits causality beyond animal data, excluding human confounders like metformin (Cuesta-Zuluaga *et al*., 2017) or diet;16S resolution misses’ strain-level metabolism captured by metagenomics (Slouha *et al*., 2023). Strengths include PRISMA-compliant screening, multi-EDC breadth, and functional grouping, establishing an EDC-diabetogenic microbiome atlas comparable to obesity benchmarks (Davis *et al*., 2016). The meta-analysis provides the distinct pattern of microbial alteration on the EDCs exposure.

## Conclusion

This meta-analysis of 16S rRNA data from 10 studies on 14 endocrine-disrupting chemicals (EDCs) demonstrates consistent gut microbiota dysbiosis linked to hyperglycaemia in type 2 diabetes models (Fig. 8). Phylum-level changes include elevated Firmicutes/Bacteroidetes ratio, increased Desulfobacterota and Patescibacteria, and depleted Bacteroidota. Genus-level shifts show depletions in beneficial taxa *(Akkermansia, Bifidobacterium, Lactobacillus, Faecalibacterium)*, enrichments in xenobiotic-degraders (Bacteroides, Bilophila, Desulfovibrio, Parabacteroides), and pro-inflammatory genera (*Alistipes, Collinsella, Escherichia-Shigella, Streptococcus*). These microbial signatures align with biochemical markers like hyperglycemia, dyslipidemia, and elevated liver enzymes (ALT, AST, ALP), supporting EDCs’ role in T2DM progression via gut-mediated pathways such as SCFA suppression, inflammation, and barrier disruption. Unlike obesity-driven dysbiosis, EDC effects uniquely promote xenobiotic metabolism and chemical toxicodynamics, distinguishing their diabetogenic profiles.

**Fig. 8.**
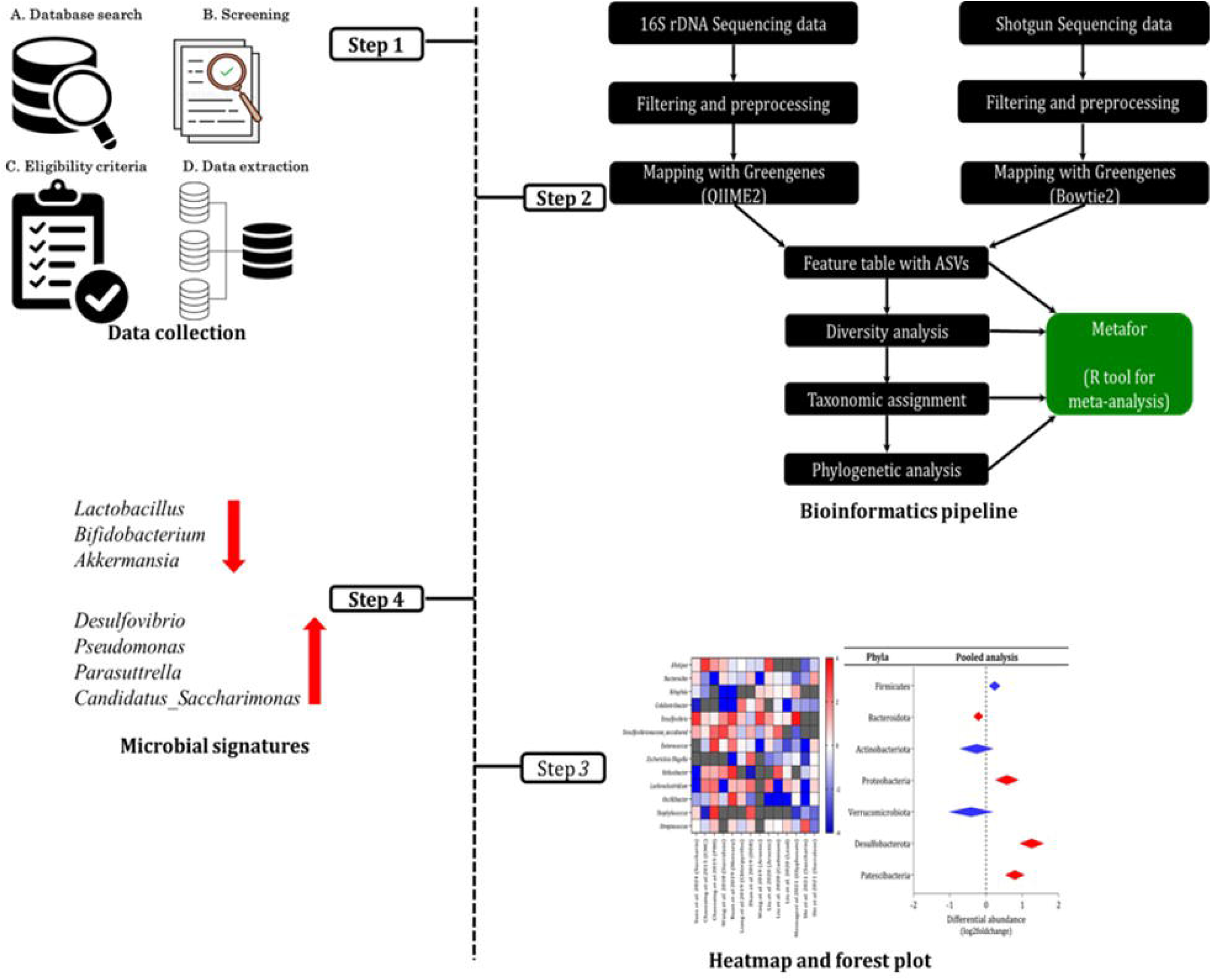
Graphical Abstract of the study.

## Abbreviations

16S rRNA: 16S Ribosomal Ribonucleic Acid
ALP: Alkaline Phosphatase
ALT: Alanine Aminotransferase
AST: Aspartate Aminotransferase
CI: Confidence Interval
DADA2: Divisive Amplicon Denoising Algorithm 2
EDCs: Endocrine-Disrupting Chemicals
F/B ratio: Firmicutes to Bacteroidetes Ratio
GLP-1: Glucagon-Like Peptide-1
HbA1c: Hemoglobin A1c (Glycated Hemoglobin)
LEfSe: Linear Discriminant Analysis Effect Size
LPS: Lipopolysaccharide
NF-κB: Nuclear Factor Kappa B
PCoA: Principal Coordinates Analysis
PICRUSt: Phylogenetic Investigation of Communities by Reconstruction of Unobserved States
PPARγ: Peroxisome Proliferator-Activated Receptor Gamma
PRISMA: Preferred Reporting Items for Systematic Reviews and Meta-Analysis
QIIME 2: Quantitative Insights into Microbial Ecology 2
SCFA: Short-Chain Fatty Acids
SMDs: Standardized Mean Differences
SRA: Sequence Read Archive
T2DM: Type 2 Diabetes Mellitus
TLR4: Toll-Like Receptor 4
TMAO: Trimethylamine N-Oxide

## Ethics approval and Consent for participation

Not applicable

## Consent for publication

Not applicable

## Author contributions

Conceived and designed the experiments: KS, MG & GV; Performed search and collected data: KD, BG, HG and GV; Analyzed the data: GM, KD, BG and GV. Wrote the manuscript: KD, BG and GV; Revised the manuscript: GM, HG, KS and MG. All authors read and approved the final manuscript.

## Acknowledgements

The authors acknowledge Founding trustees, President of KMCH Research Foundation for their support and access to all resources.

## Disclosure of potential conflicts of interest

The authors declare no potential conflicts of interest

## Data Availability Statement

All data are included in the manuscript and supplementary files

## Funding

This study is funded by Indo-French Centre for Promotion of Advanced Research (CEFIPRA) under CSRP Scheme (Project No. 6303-3). KD, BG and GM acknowledge Indian Council for Medical Research, Government of India for awarding Senior Research Fellowship (Fellowship ID: 2021-14115) and Research Associate fellowship (Fellowship ID: 2020-6690) (Fellowship ID: 2021-12337) respectively.

## Notes

### Competing Interest Statement

The authors have declared no competing interest.

