## Supplementary file for "Meta-analysis reveals a distinct and uniform gut microbial signature associated with endocrine-disrupting chemicals-induced diabetes"

**Supplementary Information:**

**Identification of common gut microbial signature in Endocrine-disrupting chemical-induced diabetes rodent models: A Meta-analysis Approach**

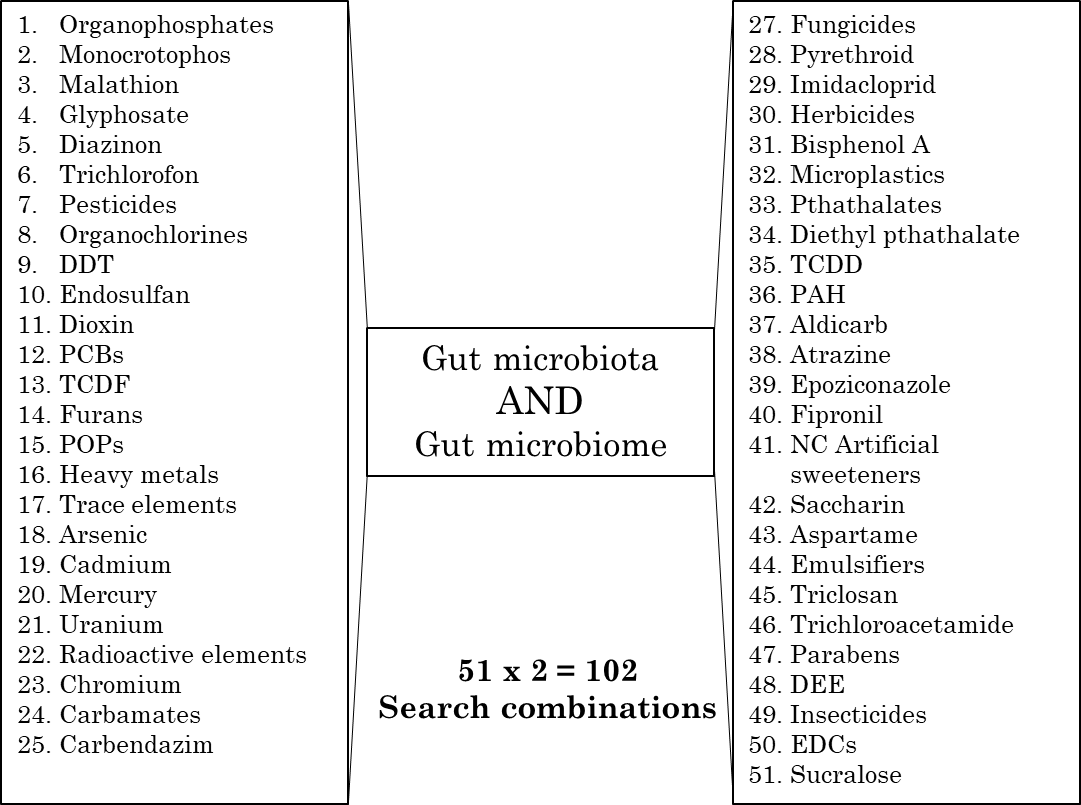

**Fig. S1: Systematic PubMed Search for Peer-Reviewed Articles: Endocrine-Disrupting Chemicals (EDCs) and Gut Microbiome/Microbiota Keywords**

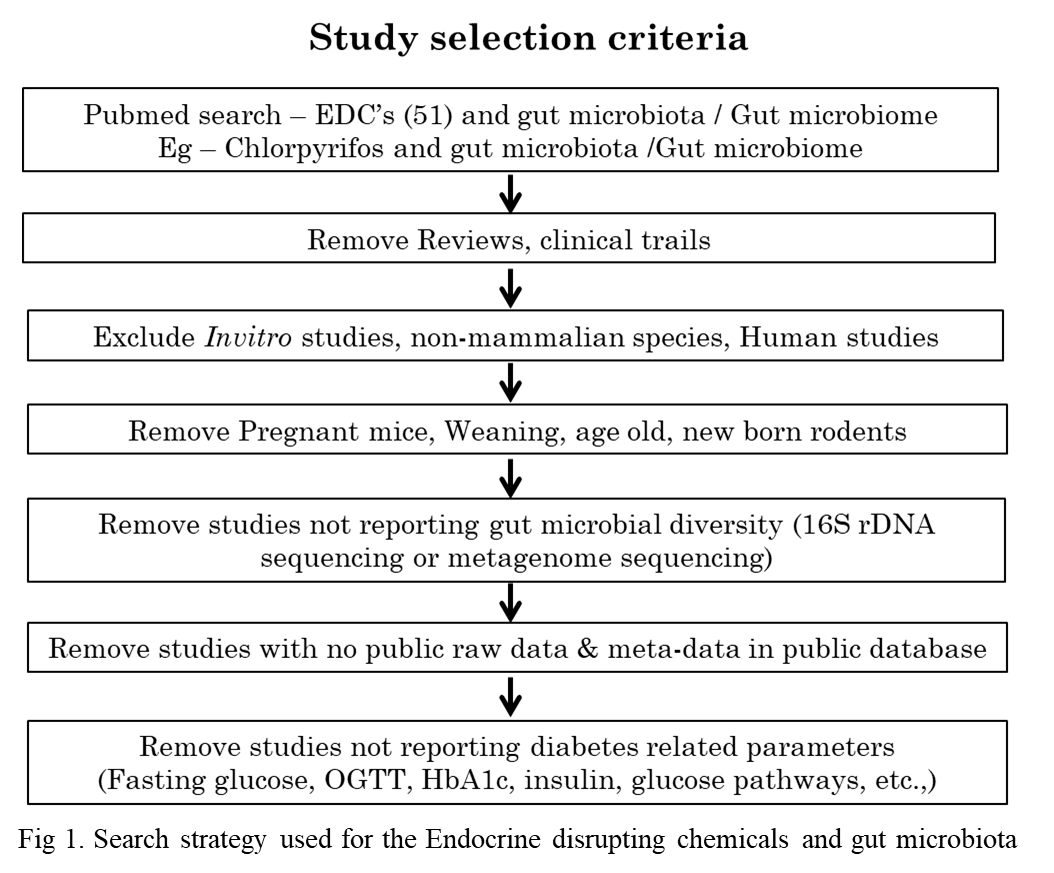

**Fig. S2: Search strategy of the PRISMA guidelines.** Used for the study selection between the endocrine disrupting chemicals and gut microbiota.

**Fig. S4:** Phylum-level microbial composition across chemical exposure categories.

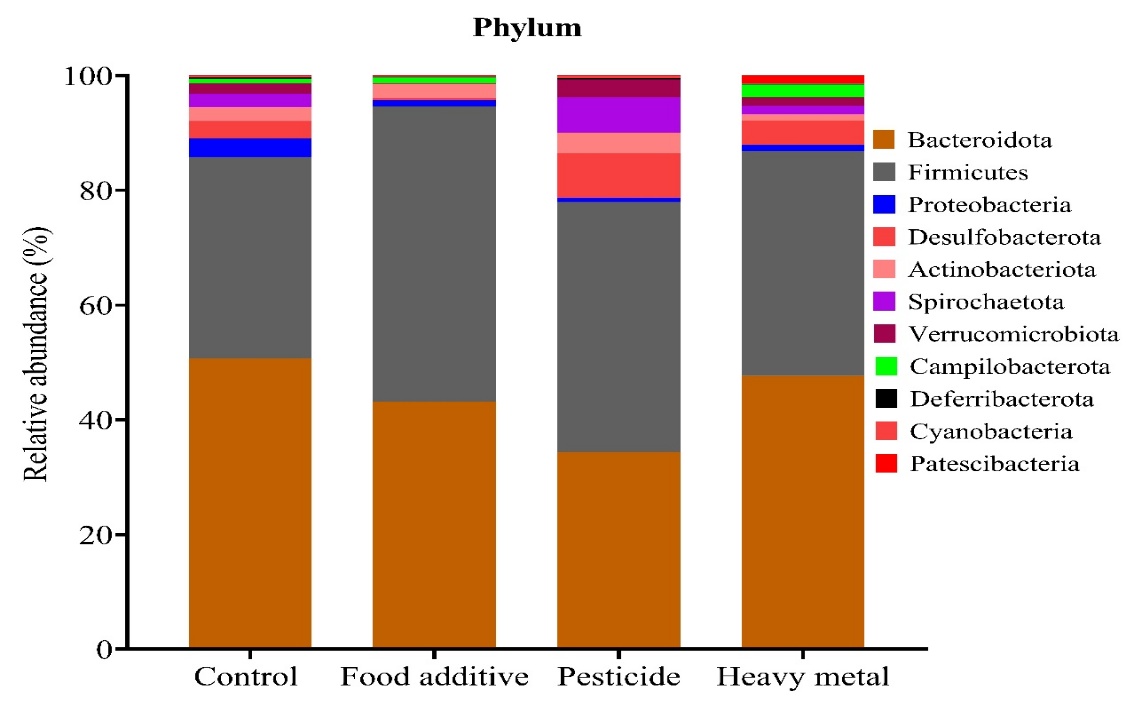

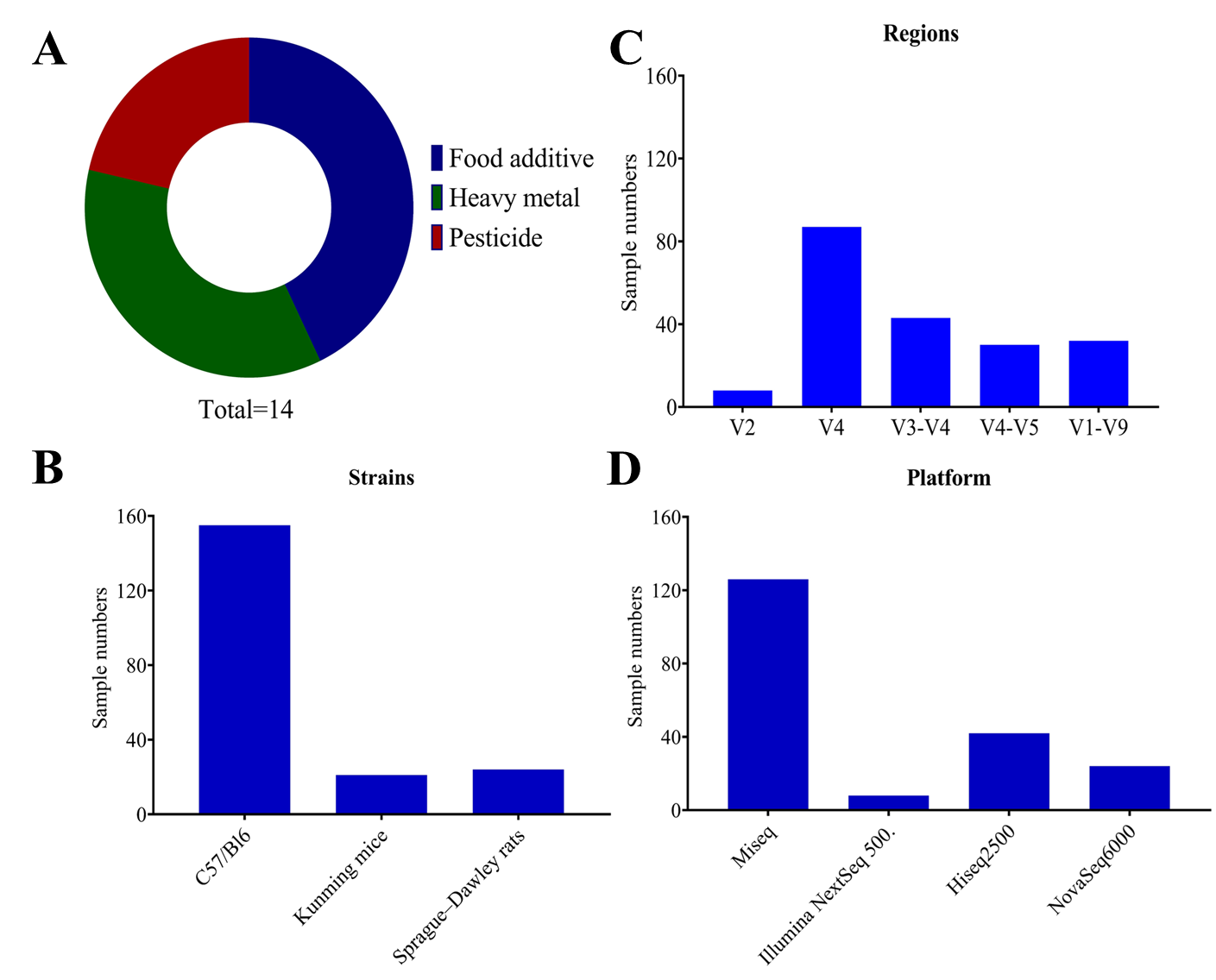

**Fig. S3:** Details of 10 selected studies. A – categorization of endocrine disrupting chemicals on selected articles. B – Different rodent strains. C- Types of the variable regions. D – Platforms used in the articles.

**Table S1**: Summary of Experimental Parameters Related to Metabolic, Hepatic, and Inflammatory Outcomes Across Publications

| **Parameters** | **Publication** | | | | | | | | | |
| --- | --- | --- | --- | --- | --- | --- | --- | --- | --- | --- |
|  | **01** | **02** | **03** | **04** | **05** | **06** | **07** | **08** | **09** | **10** |
| Body weight | ↑ | ↑ | ↓ | ↓ | ↑ | ↑ | NC |  | NC |  |
| Fat Weight |  | ↑ |  |  | ↑ | ↑ |  |  |  |  |
| Food intake | ↓ | ↑ | ↑ |  | ↓ | ↓ |  |  | ↓ |  |
| Water intake | ↑ |  | ↑ |  |  |  |  |  | ↑ |  |
| Liver gene expression |  |  |  |  |  | ✓ |  |  |  | ✓ |
| Liver lipid content |  |  |  |  |  | ✓ |  |  |  | ✓ |
| EDC accumulation |  |  |  |  |  |  |  | ✓ |  |  |
| AhR |  |  |  |  |  |  |  |  |  | ✓ |
| Tryptophan metabolites | ✓ |  |  |  |  |  |  |  |  |  |
| Liver Histopathology |  |  |  |  |  | ✓ |  |  |  | ✓ |
| Fasting insulin | ↑ | ↑ | ↑ | ↑ | ↑ | ↑ | ↑ | ↑ | ↑ | ↑ |
| Lipid profile (TC, LDL-c& TG) | ↑ | ↑ |  |  | ↑ |  |  |  |  |  |
| AST | NC |  |  |  | ↑ | ↑ |  |  |  |  |
| ALT |  |  |  |  |  | ↑ |  |  |  | ↑ |
| Uric acid |  |  |  |  |  |  |  | ↑ |  |  |
| Creatinine |  |  |  |  |  | ↑ |  | ↑ |  | ↑ |
| Total bile acids |  |  |  |  |  |  |  | ↑ |  |  |
| TNF-α, PA-1, MCP-1 |  |  |  |  |  |  |  | ↑ |  |  |
| Interleukins |  |  |  |  |  | ↑ |  |  |  | ↑ |
| CD-14 |  |  |  |  | ↑ |  |  |  |  |  |
| ↑ - increased, ↓ - decreased, NC – No change, ✓ - Observed | | | | | | | | | | |

**Table S2:** Summary of sequencing details and accession number for the selected articles

| **S.No.** | **Publication Details** | **16S/Shotgun sequencing** | **Reads** | **Database** | **Accession Number** | **No. of samples** |
| --- | --- | --- | --- | --- | --- | --- |
| 01 | Suez *et al.* 2014.  *Nature* | Shotgun sequencing | Single end | ENA | PRJEB6996 | 8 |
| 02 | Chassaing *et al.* 2015. *Nature* | 16S rDNA Sequencing | Paired end | ENA | PRJEB8035 | 66 |
| 03 | Wang QP *et al.* 2018. *PLoS One* | 16S rDNA Sequencing | Paired end | NCBI | SRP148650 | 8 |
| 04 | Ruan *et al.* 2019. *Biol. Trace Elem. Res.* | 16S rDNA Sequencing | Paired end | NCBI | PRJNA418396 | 10 |
| 05 | Liang *et al.* 2019. *Microbiome* | 16S rDNA Sequencing | Paired end | NCBI | PRJNA510290 | 12 |
| 06 | Zhan *et al.* 2019.  *Environ. Int.* | 16S rDNA Sequencing | Paired end | NCBI | PRJNA529558 | 10 |
| 07 | Wang *et al.* 2019. *Environ. Int.* | 16S rDNA Sequencing | Paired end | NCBI | PRJNA552267 | 18  F**ig. 5.1.10: Differentially abundant gut bacterial taxa identified by LEfSe analysis.** Cladogram showing the taxonomic hierarchy of gut bacterial genera that differ significantly between study groups, as identified by LEfSe analysis (LDA score above the significance threshold). Colored branches and nodes represent taxa enriched in each group, while black nodes indicate taxa not significantly differentially abundant. |
| 08 | Liu *et al.* 2020. *Front. Microbiol.* | 16S rDNA Sequencing | Paired end | NCBI | PRJNA596575 | 24 |
| 09 | Mesnage *et al.* 2021. *Environ. Health Perspect.* | Shotgun sequencing | Paired end | NCBI | PRJNA609596 | 24 |
| 10 | Shi *et al.* 2021. *mSystems* | Shotgun sequencing | Paired end | NCBI | PRJNA631099 | 9 |
